# Trends in Pacific Canadian groundfish stock status

**DOI:** 10.1101/2021.12.13.472502

**Authors:** Sean C. Anderson, Brendan M. Connors, Philina A. English, Robyn E. Forrest, Rowan Haigh, Kendra R. Holt

## Abstract

We assembled estimated biomass (B) time series from stock assessments for 24 Pacific Canadian groundfish stocks and modelled average and stock status through 2020 based on biomass relative to each stock’s (1) Limit Reference Point (B/LRP), (2) Upper Stock Reference (B/USR), and (3) biomass at maximum sustainable yield (B/B_MSY_). The overall mean B/LRP in 2020 was 3.2 (95% credible interval [CI]: 2.6–3.9). The overall mean B/USR and B/B_MSY_ in 2020 was 1.5 (95% CI: 1.3–1.9) and 1.4 (95% CI: 1.1–1.7), respectively. Average stock status declined from 1950 to around 2000 and has remained relatively stable since then. The change around 2000 followed the implementation of ITQs (individual transferable quotas) for the trawl fleet and the commencement of the synoptic trawl surveys. As of their last assessment, four stocks (Strait of Georgia Lingcod [Area 4B], coastwide Bocaccio, and inside and outside Quillback Rockfish) had a greater than 5% probability of being below their LRP (i.e., in the “ critical zone”); Pacific Cod in Area 3CD had a 4.6% probability. Roughly one-third of stocks had a greater than 1 in 4 chance of being below their USR (i.e., in the “ cautious zone”). Conversely, two-thirds of assessed groundfish stocks had a high (>75%) probability of being above the USR (i.e., in the “ healthy zone”).

## Introduction

Recent amendments to Canada’s *Fisheries Act* via the <u>Fish Stocks provisions</u> have increased DFO’s focus on stock status with respect to biological reference points. The amendments are based on Canada’s Precautionary Approach (PA) Framework, which describes two stock status reference points: (1) the Limit Reference Point (LRP), a “ status below which serious harm is occurring to the stock” ; and (2) the Upper Stock Reference Point (USR), which represents the “ threshold below which removals must be progressively reduced in order to avoid reaching the LRP” (DFO 2009, Figure 1a). The USR defines the breakpoint between the healthy and cautious zone and the LRP defines the breakpoint between the cautious and critical zone (Figure 1a). The LRP and USR are often calculated based on B_MSY_ (biomass at maximum sustainable yield) at provisional levels of 0.4 B_MSY_ and 0.8 B_MSY_, but in other cases can be based on proxies, such as average or minimum historical biomass levels (e.g., Forrest *et al*. 2020).

**Figure 1:**
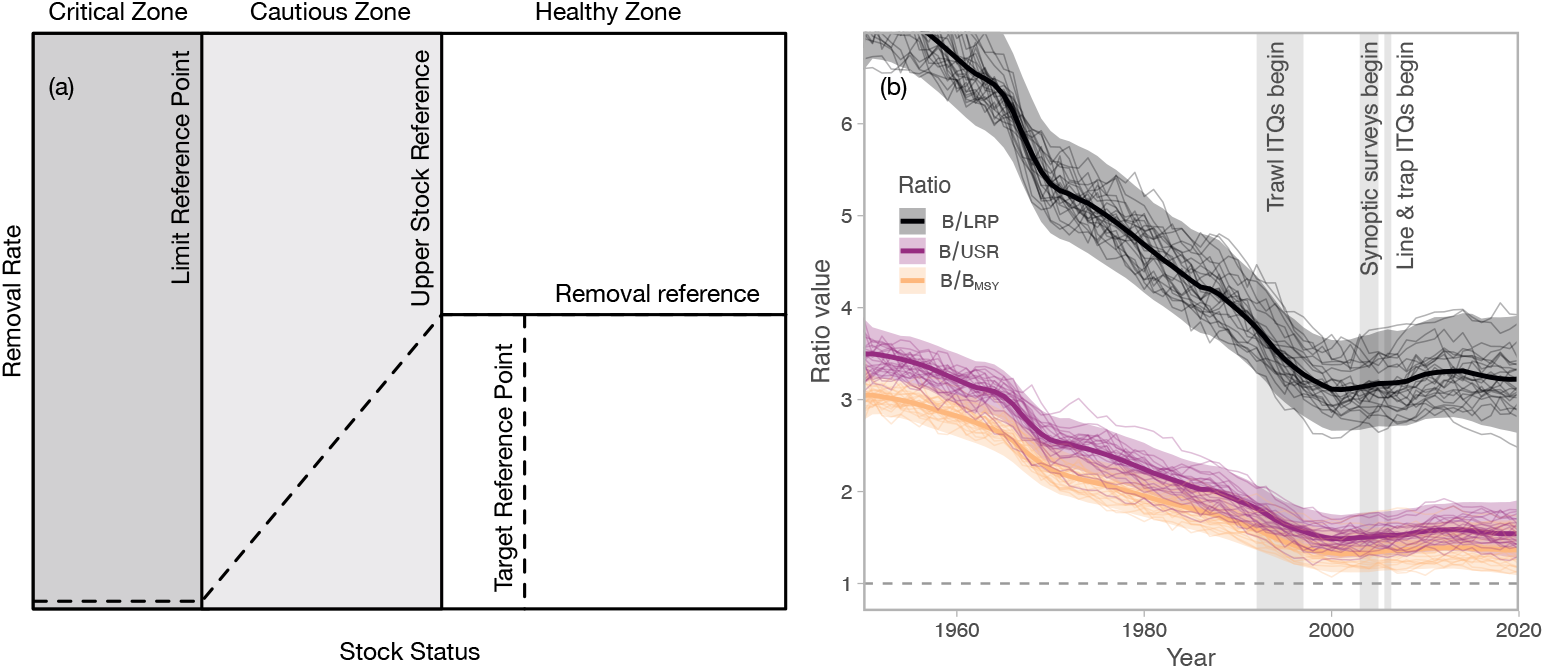
Illustration of DFO’s Precautionary Approach Framework. The two vertical lines—the Limit Reference Point (LRP) and Upper Stock Reference (USR)—are the focus of this analysis. Based on DFO (2009). (b) Overall mean biomass status (*x*_*t*_) across all stocks for B/LRP, B/USR, and B/B_MSY_. Dark lines represent the median, shaded ribbons represent 95% quantile credible intervals, and thin lines represent 25 illustrative draws from the posterior distribution.

Assessment scientists conduct regular stock assessment for major fish stocks in BC. These assessments combine fishery-dependent data (such as commercial catches) with fishery-independent data (data from scientific surveys) to estimate quantities such as spawning stock biomass, growth and maturity, and derive measures of stock status and fishing intensity. Here, we gathered output data from assessments for 24 stocks representing the Bayesian posterior distribution of estimated biomass (B) compared to three measures of stock status: (1)

B/LRP (biomass divided by the LRP), (2) B/USR (biomass divided by the USR), (3) B/B_MSY_ (biomass divided by B_MSY_). We developed a hierarchical Bayesian state-space time-series model to explore trends in these measures of status across all stocks until the year 2020. To do this, we built on a model described in Hilborn *et al*. (2020) in two ways: (1) we incorporated uncertainty in stock-specific status; and (2) we projected the latent underlying stock-specific status forward to the last year (2020). The latter accounts for not all stocks having an up-to-date stock assessment and accounts for future possible stock states based on the time-series properties of the fitted stock trajectories.

## Methods

We modelled overall (i.e., all stocks combined) log stock status *x* at time *t* as *x*_*t*=1_ = *x*_0_ and 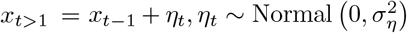, where *η*_*t*_ is a random walk deviation with standard deviation*σ*_*η*_. We assumed an auto-regressive observation model:

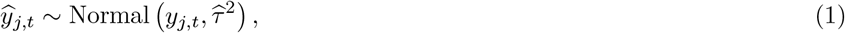

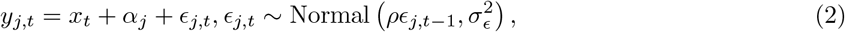

where 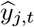 is the mean log stock status for stock *j* and time *t* (from an assessment), *y j,t* is the unobserved “ true” mean stock-specific status, and 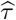 is the standard deviation of the log stock status from an assessment. The symbol *α*_*j*_ represents a stock-specific intercept that is constrained such that the sum of all *α*_*j*_ is zero to make the overall mean *x*_*t*_ identifiable. The symbol *ϵ*_*j,t*_ represents a first-order autoregressive (AR1) deviation with correlation *ρ* and standard deviation*σ*_*ϵ*_. We placed half-Normal(0, 1) priors on*σ*_*ϵ*_ and*σ*_*η*_, Normal(0, 5^2^) priors on *x*_0_ and *α*_*j*_, and a Normal(0, 1)[-1, 1] prior on *ρ*. We fit the models with Stan (Carpenter *et al*. 2017; Stan Development Team 2020), sampling 1000 iterations on each of 6 chains and discarding the first half as warm up. Data and code to reproduce our analysis are available at https://github.com/pbs-assess/gftrends with the specific version at https://github.com/pbs-assess/gftrends/releases/tag/v0.01.

## Results

Of the 24 groundfish stock status time series included in the analysis (Table 1), four stocks provided historical reference points for management advice, 18 provided MSY-based reference points, and two (Rocksole in Areas 5AB and 5CD) provided both historical and MSY reference points (we herein use the MSY-based ones). The most recent assessments were for the Rougheye/Blackspotted Rockfish complex, Pacific Cod, Bocaccio, inside and outside Yelloweye Rockfish, and Sablefish.

**Table 1:** Groundfish stocks included in the analysis with associated data reference. In cases where a Research Document was still ‘In press’, we indicate the associated published Science Advisory Report or Science Response.

| Stock | Reference |
| --- | --- |
| Arrowtooth Flounder BC | Grandin & Forrest (2017) |
| Bocaccio BC | DFO (2020a) |
| Lingcod 4B | Holt *et al.* (2016a) |
| Pacific Ocean Perch 3CD | Edwards *et al.* (2014) |
| Pacific Ocean Perch 5ABC | Edwards *et al.* (2013) |
| Pacific Ocean Perch 5DE | Edwards *et al.* (2011) |
| Pacific Cod 3CD | Forrest *et al.* (2020) |
| Pacific Cod 5ABCD | Forrest *et al.* (2020) |
| Quillback BC Outside | Yamanaka *et al.* (2011) |
| Quillback VI Inside | Yamanaka *et al.* (2011) |
| Redstripe Rockfish BC N | Starr & Haigh (2021a) |
| Redstripe Rockfish BC S | Starr & Haigh (2021a) |
| Rocksole 5AB | Holt *et al.* (2016b) |
| Rocksole 5CD | Holt *et al.* (2016b) |
| Rougheye/Blacksp. BC N | DFO (2020d) |
| Rougheye/Blacksp. BC S | DFO (2020d) |
| Sablefish BC | DFO (2020b) |
| Shortspine Thornyhead BC | Starr & Haigh (2017) |
| Silvergray Rockfish BC | Starr *et al.* (2016) |
| Walleye Pollock BC N | Starr & Haigh (2021b) |
| Walleye Pollock BC S | Starr & Haigh (2021b) |
| Widow Rockfish BC | DFO (2019b) |
| Yelloweye Rockfish 4B | DFO (2020c) |
| Yellowtail Rockfish BC | DFO (2015) |

Across all stocks, there was a decline in average stock status for all three indices prior to approximately 2000 (Figure 1b). The late 1990s and early 2000s marked the beginning of a relatively stable average status. We estimated the overall mean B/LRP in 2020 to be 3.2 (95% credible interval [CI]: 2.6–3.9). The overall mean B/USR and B/B_MSY_ in 2020 was 1.5 (95% CI: 1.3–1.9) and 1.4 (95% CI: 1.1–1.7), respectively. Uncertainty in average status in the last few years increased because not all stocks had a current assessment (Figure 1b).

The clear pattern in the average biological status has been accompanied by considerable variation within and across individual stocks (Figure 2). Within individual stocks, Pacific Cod and Walleye Pollock—both with shorter generation times compared to many other stocks here—exhibited strong patterns of decadal variation. Across stocks, Redstripe Rockfish in northern and southern BC and coastwide Yellowtail Rockfish are examples of stocks with B/LRP trajectories consistently above the average. Conversely, Lingcod in 4B and coastwide Bocaccio are examples of stocks with B/LRP estimated consistently below the average. Sablefish followed the average B/LRP trajectory until around 1990 when they continued to decline while the average stabilized. The Sablefish stock trajectory has shown signs of increase since around 2015 as a result of large recent recruitments (DFO 2020b, Figure 2).

**Figure 2:**
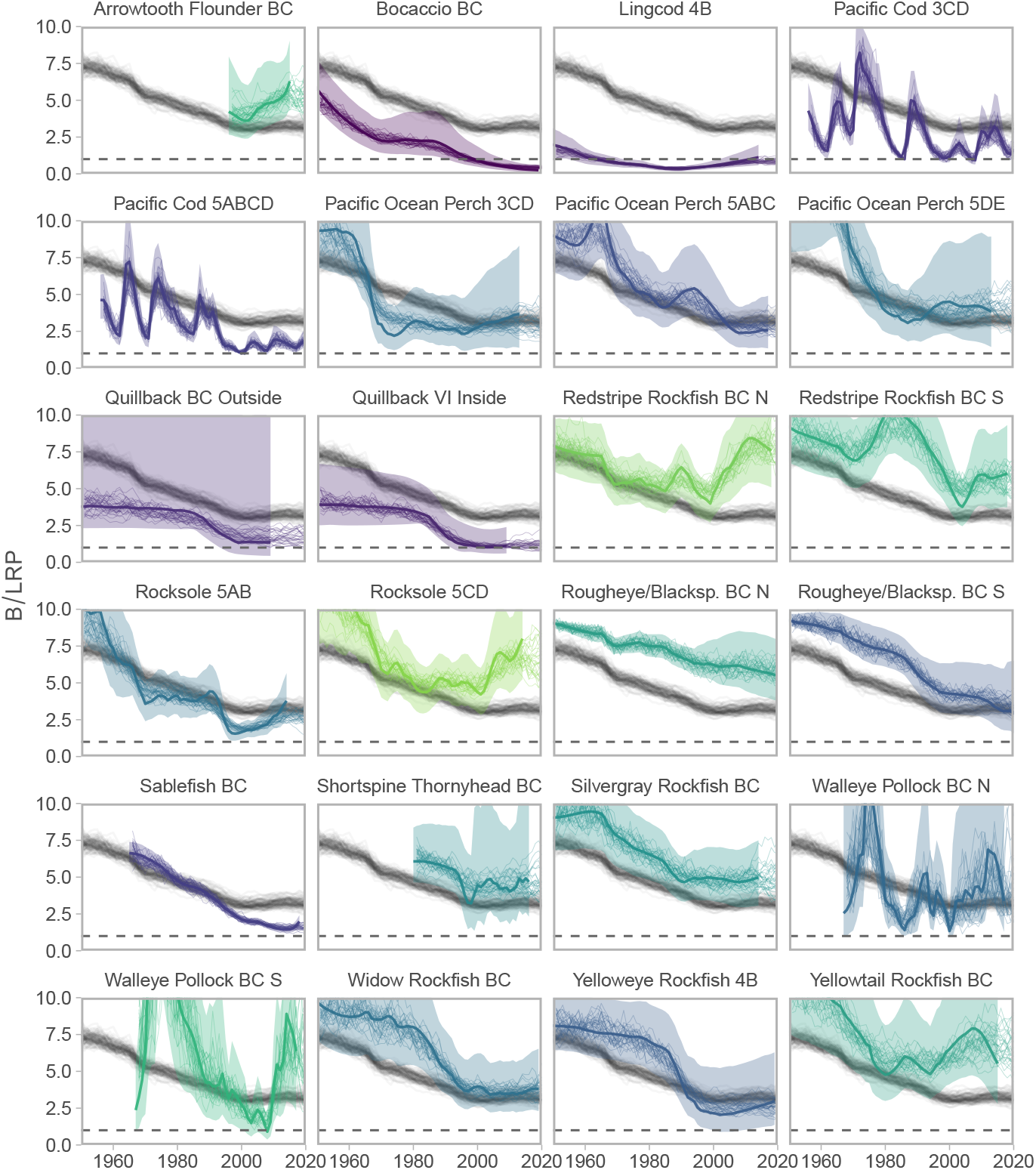
Trends in B/LRP for 24 groundfish stocks in BC. Coloured lines and ribbons represent the individual stocks. Colours represent the ratio in the last year of assessment such that green stocks are highest and purple are lowest. Dark coloured lines and shaded ribbons represent output from stock assessments: trajectories of median B/LRP 95% quantile credible intervals. Thin lines represent draws from the posterior distribution of *y*_*j,t*_ (latent stock-specific mean trends; colours) and *x*_*t*_ (latent overall mean trend; grey). The overall mean trend (*x*_*t*_; grey) is the same across panels (Figure 1b).

Estimated biomass was above the LRP and USR for most stocks as of the most recent assessment (Figure 3). Lingcod in 4B and coastwide Bocaccio were the only stocks (with full posterior distributions available) with *>* 5% posterior density below their LRP as of their most recent assessment (Figure 3). Pacific Cod in 3CD almost met this threshold with a 4.6% probability B *<* LRP in 2020. The assessment of Bocaccio, however, projected the stock to rebuild above the LRP by 2023 due to a very large recruitment event in 2016 (DFO 2020a). Quillback Rockfish in both outside and inside waters had *>* 5% posterior density below their LRPs as of the 2011 assessment, but the full posterior distribution was not available (quantiles are shown in Figure 2). Considering the USR instead of the LRP, 7/22 of the stocks in Figure 3 had > 25% probability of being below their USR as of their most recent assessment.

**Figure 3:**
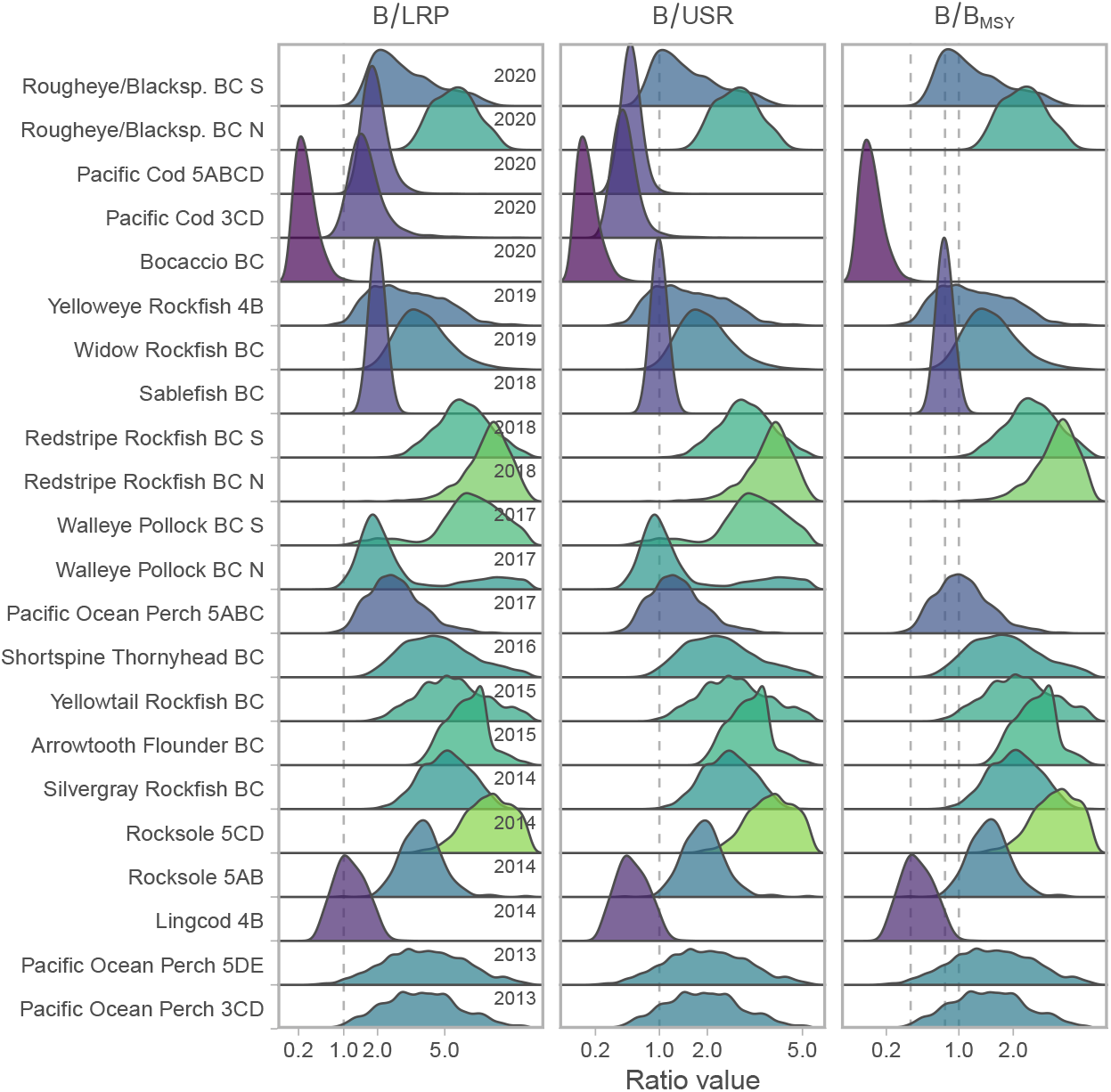
Posterior distribution of the three measures of stock status for 22 of the 24 stocks. Stocks are arranged in order of assessment with the most recent assessments at the top; years in the first column represent the year the status represents. Colours represent the mean B/LRP value such that green is highest and purple is lowest. Stocks with missing data in the B/B_MSY_ column are assessments where historical reference points were used instead of MSY-based reference points. Vertical dashed lines are at values of 1.0 in all columns and also at values of 0.4 and 0.8 in the B/B_MSY_ column (0.4 and 0.8 B_MSY_ are provisional LRP and USR values in the PA Framework). The x-axis has been square-root transformed to slightly compress high ratio values for visualization. In a few cases (e.g., Bocaccio) the status represents a one- or two-year projection from the last year of available data. The full posteriors were not available for Quillback Rockfish stocks, which are shown in Figure 2.

## Discussion

The factors most likely influencing the trends illustrated here have been fishery removals and management interventions. The transition from declining average B/LRP and B/USR to a relatively stable trajectory coincided with (1) the implementation of individual transferable quotas (ITQs) for the trawl fleet, (2) the introduction of 100% at-sea observer coverage over the period 1992–1997 (Turris 2000), and (3) the initiation of the current synoptic trawl surveys in 2003. Furthermore, ITQs and electronic at-sea monitoring were introduced into the longline and trap fisheries in 2006 (Stanley *et al*. 2015). Following these major management changes, quota for many stocks has remained relatively constant over the last two decades (DFO 2019a). The decadal patterns in assessed status for Pacific Cod (Forrest *et al*. 2020) and Walleye Pollock (Starr & Haigh 2021b) in early decades are largely driven by trends in commercial catches and catch per unit effort. Consistently poor recruitment for decades is thought to be the primary driver behind stock declines for some rockfish such as Bocaccio (DFO 2020a), although many stocks are data-limited and there is therefore considerable uncertainty around recruitment trends, which may be confounded with other factors such as the management changes noted above. In terms of absolute stock status, estimates of B_MSY_ for data-limited species may be highly uncertain, or in some cases biased (Forrest *et al*. 2018).

There are also potential species interactions and climatic effects on trends in local biomass density of these species. Loss of spawning habitat due to recent oceanographic conditions has been hypothesized to have led to years of low recruitment in some groundfish (e.g., Pacific Cod in nearby Alaskan waters, Laurel & Rogers 2020). Conversely, after decades of poor recruitment, Bocaccio experienced a year of extremely strong recruitment (44 times the average for Bocaccio in 2016, DFO 2020a) possibly due to the availability of oxygen-rich water at depth during gestation (Schroeder *et al*. 2019). The slow recovery of Area 4B Lingcod since initial fishery-driven declines between the 1930s and 1980s may be a result of changes in the Strait of Georgia ecosystem (e.g., plankton timing, pinnipeds, Holt *et al*. 2016a). Underlying these stock-level trends, recent work suggests that temperature velocity—the pace a fish would have to move to maintain consistent temperature—may be related to a fine-scale redistribution of groundfish species density in Canadian Pacific waters (English *et al*. 2021).

The Sustainable Fisheries Framework and the Fish Stocks provisions of the *Fisheries Act* require that major fish stocks be maintained above their LRP with high probability. Stocks below their LRP will require a rebuilding plan. While average stock status was clearly above the LRP as of 2020, there was considerable variation among stocks. Four stocks had *>* 5% probability of being below their LRP and roughly one-third of assessed stocks had *>* 25% probability of being in the cautious zone where removals should be progressively reduced to avoid reaching the LRP. Rebuilding and precautionary management of stocks in the critical and cautious zones, respectively, should help ensure stock status improves over time in response to reduced fishing pressure and favourable environmental conditions if and when they occur.

